# Urbanisation and Habitat Shape Resource-Driven Dietary Shifts in Wild Birds

**DOI:** 10.1101/2025.01.17.633593

**Authors:** Marion Chatelain, Oskar Rennstam Rubbmark, Johannes Rüdisser, Michael Traugott

## Abstract

Urbanisation affects bird ecology and evolution, with changes in nutritional intake considered a key driver. However, most studies provide only snapshots of urban bird feeding ecology due to methodological limitations in analysing diets across space and time. Here, we address this gap by examining the diets of two common species with differing feeding ecologies: the great tit (*Parus major*) and blue tit (*Cyanistes caeruleus*). Diet samples from 370 birds, captured over a year at 147 locations across an urbanised landscape in Innsbruck, Austria, were molecularly analysed. Results revealed species-and season-specific diet patterns influenced by urbanisation: urban great tits exhibited higher diet diversity but reduced arthropod consumption during the breeding season, while urban blue tits compensated for lower moth intake by increasing their consumption of crab spiders and aphids. Prey consumption mirrored prey availability, highlighting resource-driven dietary shifts. Habitat type also played a significant role, with urban green spaces enhancing plant-based food diversity and residential areas increasing anthropogenic food consumption. These findings support the hypothesis that diet drives fitness and phenotypic differences between urban and rural bird populations. They highlight the need to consider both urbanisation levels and habitat characteristics to fully understand its ecological and evolutionary impacts.

## 1. Introduction

The number of urban agglomerations worldwide nearly doubled from 1990 to 2018, covering up to 24% of the land surface in some European countries (*Land Cover Data 2018*, 2022; *World Urbanization Prospects*, 2019), significantly altering abiotic conditions and land cover (Grimm et al., 2008; Lambin et al., 2001). Urban green spaces are often small, fragmented, and dominated by non-native plants (Duguay et al., 2007; Grimm et al., 2008; McKinney, 2008), reducing arthropod diversity and altering species assemblages (Chatelain et al., 2023; Fenoglio et al., 2020; Piano et al., 2020). For example, arthropod richness in urban trees declines with urbanisation, especially affecting wingless taxa such as spiders, while aphids, barklice, and dipterans become more abundant (Chatelain et al., 2023). This contrasts sharply with natural habitats and is likely a key driver of differences in bird phenotypes and fitness, as it alters food availability and quality (Isaksson, 2018; Leveau, 2021; Meyrier et al., 2017; Seress & Liker, 2015). Urban birds also rely on anthropogenic food sources provided by bird feeders and food waste (Lepczyk et al., 2004; Newsome & Rodger, 2008; Oro et al., 2013), further modifying their diets (Marzluff & Neatherlin, 2006). However, little research on urban bird diets hinders identifying the underlying mechanisms driving urbanisation-related effects on bird ecology and evolution.

Opportunistic species may, for example, adjust their diet based on available food, which can affect their nutritional intake (Robb et al., 2008), while specialists maintain consistent food preferences, which can increase their foraging effort and impact their fitness. Whatever the strategy, shifts in food availability can cascade, effecting energy allocation and reproductive success. So far, studies have reported that urban birds face lower reproductive success, likely as a result of nutrient deficiencies during the breeding season (Bailly et al., 2016; Biard et al., 2017; Chamberlain et al., 2009; Corsini et al., 2020; de Satgé et al., 2019; Heiss et al., 2009; Marciniak et al., 2007; Meillère et al., 2017; Pollock et al., 2017; Seress et al., 2018; Sinkovics et al., 2021). For example, supplementing Western jackdaw diets during egg formation resulted in larger eggs and higher hatching success (Meyrier et al., 2017). Similarly, smaller skeletal sizes (Caizergues et al., 2021; Meillère et al., 2015) and duller plumage (Leveau, 2021; Salmón et al., 2023) in urban birds have been linked to nutrient limitations. However, changes in adult bird diets in response to urbanisation was only suggested indirectly. Some studies indicate that urban birds may occupy narrower foraging niches (Peneaux et al., 2021) or have lower levels of omega-3 fatty acids in winter (Andersson et al., 2015; Isaksson et al., 2017), suggesting a reduced intake of arthropods (Isaksson et al., 2015). Despite these insights, there is a critical need for direct measurement and characterisation of adult bird diets in urban environments. This gap is largely due to the limitations of current sampling designs, which often rely on a simplistic urban vs. rural comparison, and the lack of non-invasive, sensitive methods capable of assessing diets across large numbers of individuals.

DNA-based diet analysis offers a non-invasive method for studying bird diets, enabling researchers to track changes in feeding behaviour over time and across environments (Deagle et al., 2023). This study combines two metabarcoding approaches to investigate how the diets of great tits (*Parus major*) and blue tits (*Cyanistes caeruleus*) vary across 147 locations in Innsbruck, Austria, over different seasons. Both species are common along the rural-urban gradient (Chatelain et al., 2021) and frequently visit bird feeders (Chamberlain et al., 2005; Tryjanowski et al., 2015), making them ideal for studying urbanisation and food provisioning. While both are omnivores, blue tits consume significantly more arthropods. By examining these species, we aim to capture a broader range of responses to urbanisation, addressing three key questions: (1) how does prey diversity change along the urbanisation gradient, (2) how urbanisation impacts diet composition, and (3) how human-provided food influences diets over time and space.

## 2. Methods

### 2.1. Study sites

The study was carried out within a 56.5 km² area including the populated area of Innsbruck (Austria) and its surroundings (approximately 47°13N, 11°19E – 47°17N, 11°26E; elevation ranged from 574 to 1024 m; Supplementary Figure 1). With its 104.9 km² and 131 500 habitants, Innsbruck is a small to medium size city, as the majority of cities in Europe (Nabielek et al., 2016). The city is a mosaic of commercial, residential and industrial areas, as well as few urban parks (e.g., Innsbruck Hofgarten, Rapoldi Park) and multiple small garden squares. Its woody vegetation is diverse and includes maples, beeches, birches, spruces, pines, plane trees, oaks, walnut trees, horse chestnuts, poplars, dogwoods, apple trees, sherry trees, plum trees, thuyas, *etc.* Innsbruck is surrounded by villages (e.g., Axams, Natters, Aldrans), managed coniferous and mixed forests (dominated by pines, larches, firs and beech), agricultural land and natural areas, including the 727 km² Karwendel Natural Park. There characteristics result in a pronounced gradient from urban to natural or near-natural landscapes. Such a patchy landscape offers an ideal research site to understand urban ecosystems. The 56.5 km² grid was divided into 320 cells of 500 x 500 m. Each cell centroid was considered as a potential sampling site. Out of these 320 potential sites, 180 were finally selected in a way to (i) cover the urbanisation gradient range, and (ii) assure sampling feasibility (sites located on the airport ground, in large agricultural or concrete areas without trees were excluded).

### 2.2. Urbanisation level quantification & Habitat characterisation

We estimated the percent area of distinct land use/land cover (LULC) classes within 100 and 1000 m radii around each sampling site. LULC data was derived from the Land Information System Austria (LISA) which provides a detailed vector-based dataset for the study area with 13 LC classes (Banko et al., 2014). From these data, we computed four indexes of urbanisation levels: percent of impervious surface cover and tree cover within a 100 and 1000 m radius. In addition, the habitat at each sampling site was categorized as either forest (land spanning more than 0.5 hectares with trees higher than 5 meters and a canopy cover of more than 10%; in our study area, forests were mainly composed of coniferous trees, or coniferous trees and a minority of deciduous trees), forest remnant (forest smaller than 0.5 hectares, surrounded by residential area and/or fields; in our study area, forest remnants were composed of deciduous trees, or deciduous trees and a minority of coniferous trees), residential area, garden square (small green space with recreational purpose), park (large green space with recreational purpose) or business park (industrial and commercial area). Habitats were classified independently of urbanisation level. For example, residential areas could be located within the city of Innsbruck or in a neighbouring rural village.

### 2.3. Bird sampling

In total, 222 Great tits (*Parus major*) and 148 Blue tits (*Cyanistes caeruleus*) were sampled at 147 out of the 180 sampling sites. Birds were caught every other month from October 2020 to August 2021. Each sampling campaign lasted 10 days and birds were caught at three different sites per day, meaning that 30 locations were sampled per sampling campaign. The sites were selected to cover the rural-urban gradient in each sampling campaign. This objective was largely achieved, with the median, minimum and maximum imperviousness within a 100 m radius ranging from 17% to 25%, 0 to 1%, and 68% to 94%, respectively. Birds were attracted to a mist-net using a loud-speaker playing calls from the two focal species. After taking a series of morphometric measurements, a dropping was collected by placing the bird in a purpose-built box (i.e., an opaque PVC box equipped with perches separated with a grid from a DNA-free compartment at the bottom where the dropping falls), kept in a cool box before being frozen at-80°C.

### 2.4. Molecular diet analysis

Whole droppings were homogenized with 410 µl lysis buffer containing 10 µl Proteinase K (10 mg/mL, AppliChem, Darmstadt, Germany) and TES-buffer (0.1 M TRIS, 10 mM EDTA, 2% SDS, pH 8) and was incubated at 56° overnight. The DNA was extracted from 200 µl of lysate on a BioSprint96 automatic extraction platform using the BioSprint96 DNA Blood Kit (Qiagen, Hilden, Germany). Four negative extraction controls (DNA extraction blanks) were included to monitor for carry-over DNA contamination during the extraction process and were subsequently tested in PCR reactions for NGS. To detect both plant-based food and arthropod prey consumed by great tits and blue tits, we run separated PCRs with either an universal plant primer set amplifying a ∼180 to 380 bp region of the ITS2 gene: a modified version of UniPlantF2 (5’-AGGGCACGYCTGYBTGG-3’) (Guenay-Greunke et al., 2021) and a modified version of UniPlantR (5’-CGHYTGAYYTGRGGTCDC- 3’) (Moorhouse-Gann et al., 2018); or an universal arthropod primer set amplifying a ∼200 bp region of the eukaryotic CO1 gene: fwhF2 (5’-GDACWGGWTGAACWGTWTAYCCHCC- 3’) and fwhR2n (5’-GTRATWGCHCCDGCTARWACWGG- 3’) (Vamos et al., 2017). We used a nested metabarcoding approach (Daouti et al., 2024; Kitson et al., 2019). Each plate included two negative controls (molecular grade water) and two positive controls (i.e., pooled DNA from *Dionaea muscipula, Pittiosporum angustifolium* and *Honckenya peploides* for PCRs on the ITS gene, and *Agriotes sordidus, Rhipicephalus appendiculatus,* and *Cimex lectularius* for PCRs on the CO1 gene). DNA concentrations were quantified using a QIAxcel Advanced System (Qiagen, Hilden, Germany). HTS-libraries were equimolarly pooled and sequenced at the Vienna Biocenter Core Facilities VBCF (Vienna, Austria) on a NovaSeq 6000 System platform, using two lanes of the flow cell type “SP”, which allows paired-end sequencing of 250 bp.

### 2.5. Bioinformatics

Fastq reads were combined from both lanes and paired-end reads were merged using ‘vsearch’ (Rognes et al., 2016) (v2.13.4). Samples were demultiplexed allowing no fuzz (100% accuracy of forward and reverse index barcodes). Then, primers and indices were removed with ‘cutadapt’ (Martin, 2011) (v1.18). Singletons and short reads were filtered with ‘usearch’ (Edgar, 2010) (v11.0.667). Sequences were blasted with ncbi-blast/2.8.1+ (Camacho et al., 2009) against the full NCBI database. Taxonomic information was annotated to accession numbers from blast using the R-package “taxize” (Chamberlain & Szöcs, 2013) in R statistical software v 4.1.2 (R Core Team, 2022). The final dataset is based on reads with a length of at least 100 bp and a reference database match of 95% identity or higher. Due to the high risk of species misidentification when the percentage match is below 100%, diet was assessed at the order, family, and genus levels. All sequences coding for organisms other than plants or arthropods were removed. Overall, the diet of 370 individuals (148 blue tits and 222 great tits) was assessed. Sample size per species, per month and per habitat type is detailed in Supplementary Table 1.

The dataset was cleaned by removing the plant family whose prevalence is likely a result of pollen contamination. Taxa believed to primarily originate from human-provided food in bird feeders—namely *Helianthus* (sunflower seeds), *Triticum* (wheat seeds), *Cannabis* (hemp seeds), *Corylus* (hazelnuts), and *Avena* (oats)—were not removed from the dataset (Supplementary Table 2).

The co-occurrence of taxa was examined to assess the potential for secondary predation. However, we did not measure any strong correlation between any of the arthropod prey and plant taxa, suggesting that our diet communities do not include secondary predations. DNA of the bird was detected in 56 samples (i.e., six from great tits and 50 from blue tits). Bird DNA represented only a small proportion of the sequences (the percentage of sequences ranged from 0.0001 and 9.85% with a median of 0.0004%).

### 2.6. Statistical analyses

Statistical analyses were performed using R software (R Core Team, 2024). Total taxonomic richness, plant taxonomic richness and arthropod taxonomic richness at the genus, family, and order levels (i.e., the number of different taxonomic groups detected in each dropping) were compared using mixed linear models with the “lmer” function from the “lme4” package (Bates et al., 2015). Richness (centred on the monthly average) was used as the response variable, and the level of urbanisation level (measured as either impervious surface cover or vegetation cover within a 100m or 1000m radius around the sampling point) served as explanatory variable. Because we expected urbanisation to have different effects on diet composition in different seasons, the month (i.e., December, February, April, June, August and October) and its interactions with the urbanisation level were added as explanatory variables. Because several individuals have been sampled per sampling location, the location was added as random intercept. Separate models were run for great tits and blue tits. Similar models were used for comparing the proportion of frequently detected taxa (i.e., detected in at least 15% of great tits or blue tits at a specific month) over the total number of taxa. For each model, we performed a backward stepwise selection using the AIC (Sakamoto et al., 1986). A Type III Wald Chi-square test Anova was used to determine the significance of retained variables in the final models. When interactions between “urbanisation level” and “month” were retained in the models, the association between the response variable and “urbanisation level” was tested for each month using the “emtrends” function from the “emmeans” package (“emtrends” performs *t*-test adjusted for multiple comparisons; Lenth, 2024).

Differences in the taxonomic composition of great tit and blue tit diet were tested using partial linear constrained ordination methods based on Hellinger distances (RDA) on presence-absence data using the “rda” function from the “vegan” package (Oksanen et al., 2024). RDAs were performed on assemblages of frequently detected taxa only. The global models included the four indexes of urbanisation level as constraining variables. When the sample size per habitat allowed (i.e., when at least three habitats had a sample size of five or more), specifically in December and October in great tits, and in February, April and October in blue tits, the habitat type was also added as conditional variable. The environmental variables that were the most important to explain the compositional changes were identified by forward selection using the “ordistep” function from “vegan”. The variance explained by the constrained ordination and by each of the components was tested by Monte Carlo permutation test (“anova” function from “vegan”). Variation partitioning was calculated using the “rdacca.hp” function from the package of the same name (Lai et al., 2022). The contribution of each selected constraining variable to the components of the ordination was fitted by linear contribution scores using the “envfit” function. The ordination diagrams were created with the “ordiplot” function from “vegan,” which coordinates were extracted for use in “ggplot2” (Wickham et al., 2016). Ordinations were performed separately for each species and month.

Pearson correlation tests were used to assess the relationships between the occurrence of arthropod families and their availability in trees and bushes at the time of sampling. This analysis was only possible for some families, since a few taxa, while detected in bird droppings, were not detected in the environment. Moreover, individuals from some taxa were not identified to the family level. Diptera was sorted into the two suborders Nematocera and Brachycera. Hymenoptera was sorted into the paraphyletic infraorder Parasitica, the suborder Symphyta and the family Formicidae. Finally, Thomisidae and Philodromidae were not distinguished from each other and were both classified as crab spiders. Therefore, the correlations were tested for Anyphaenidae, Aphididae, Araneidae, Braconidae (with the abundance of Parasitica), Caeciliusidae, Cecidomyiidae (with the abundance of Nematocera), Ceratopogonidae (with the abundance of Nematocera), Chironomidae (with the abundance of Nematocera), Cicadellidae, Clubionidae, Curculionidae, Ectopsocidae, Elateridae, Erebidae, Formicidae, Feometridae, Noctuidae, Pentatomidae, Philodromidae (with the abundance of crab spiders), Psyllidae, Scraptiidae, Tenthredinidae (with the abundance of Symphyta), Theridiidae and Tortricidae. For those taxa, the abundance was multiplied by the percentage of vegetation cover within a 100 m radius around the sampling site to estimate their availability in the local environment where the birds were caught.

## 3. Results

### 3.1. Taxa detection

We detected 547 genera (196 plants and 351 arthropods), from 199 families (64 plants and 135 arthropods), and 51 orders (32 plants and 19 arthropods) in the diet of great tits and blue tits together. Only a proportion of the genera, families, and orders were commonly consumed (i.e., detected in the diet of at least 15% of great tits or blue tits in a given month). Specifically, 12% of genera, 27% of families, and 49% of orders were frequent in the diets of the two species. For great tits, 29 genera (21 plants and 8 arthropods), 31 families (19 plants and 12 arthropods), and 20 orders (14 plants and 6 arthropods) were commonly consumed within the study area. For blue tits, 59 genera (33 plants and 26 arthropods), 46 families (21 plants and 25 arthropods), and 22 orders (15 plants and 7 arthropods) were frequently detected in their diet (Fig. 1; Supplementary Table 3).

**Fig. 1:**
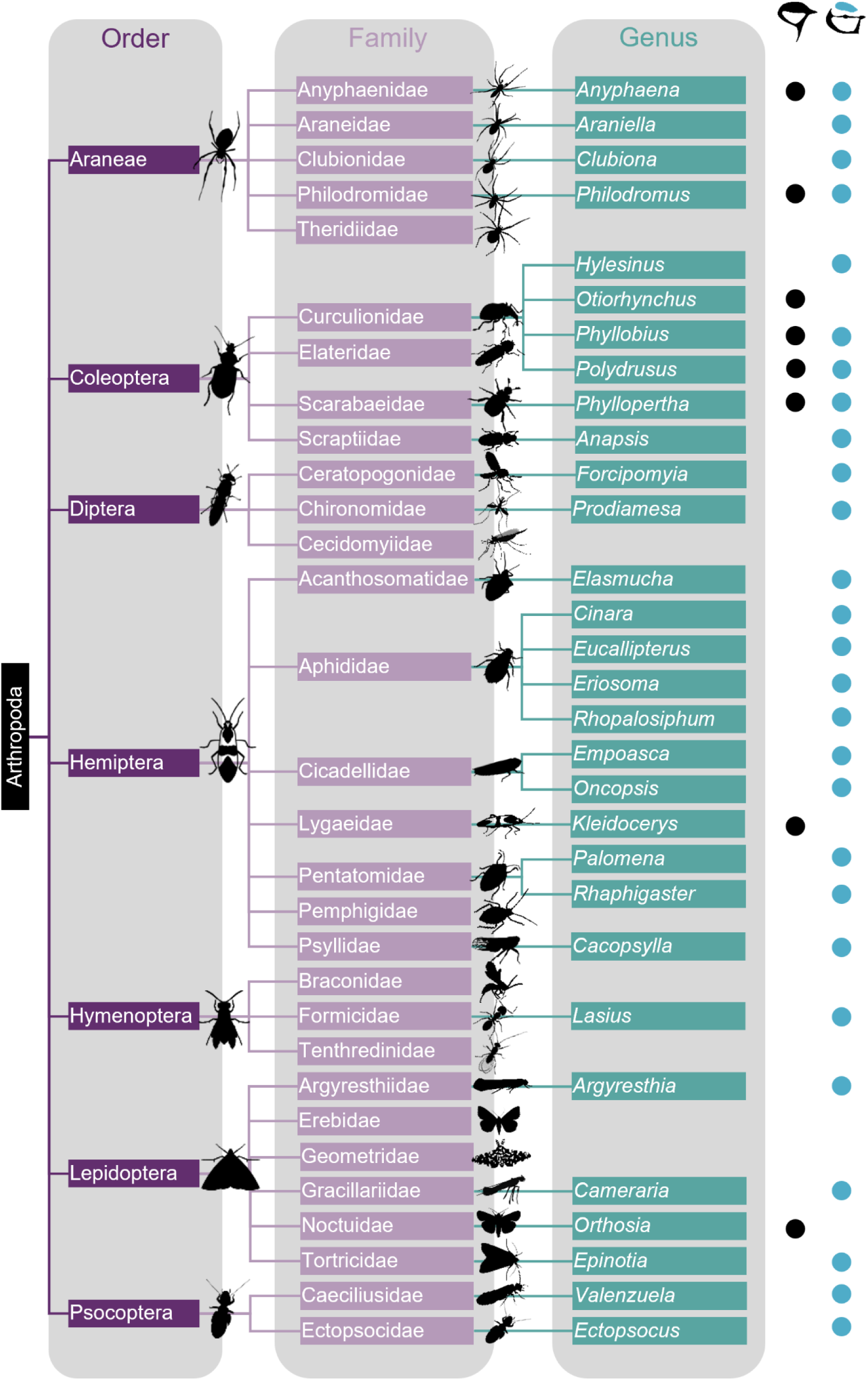
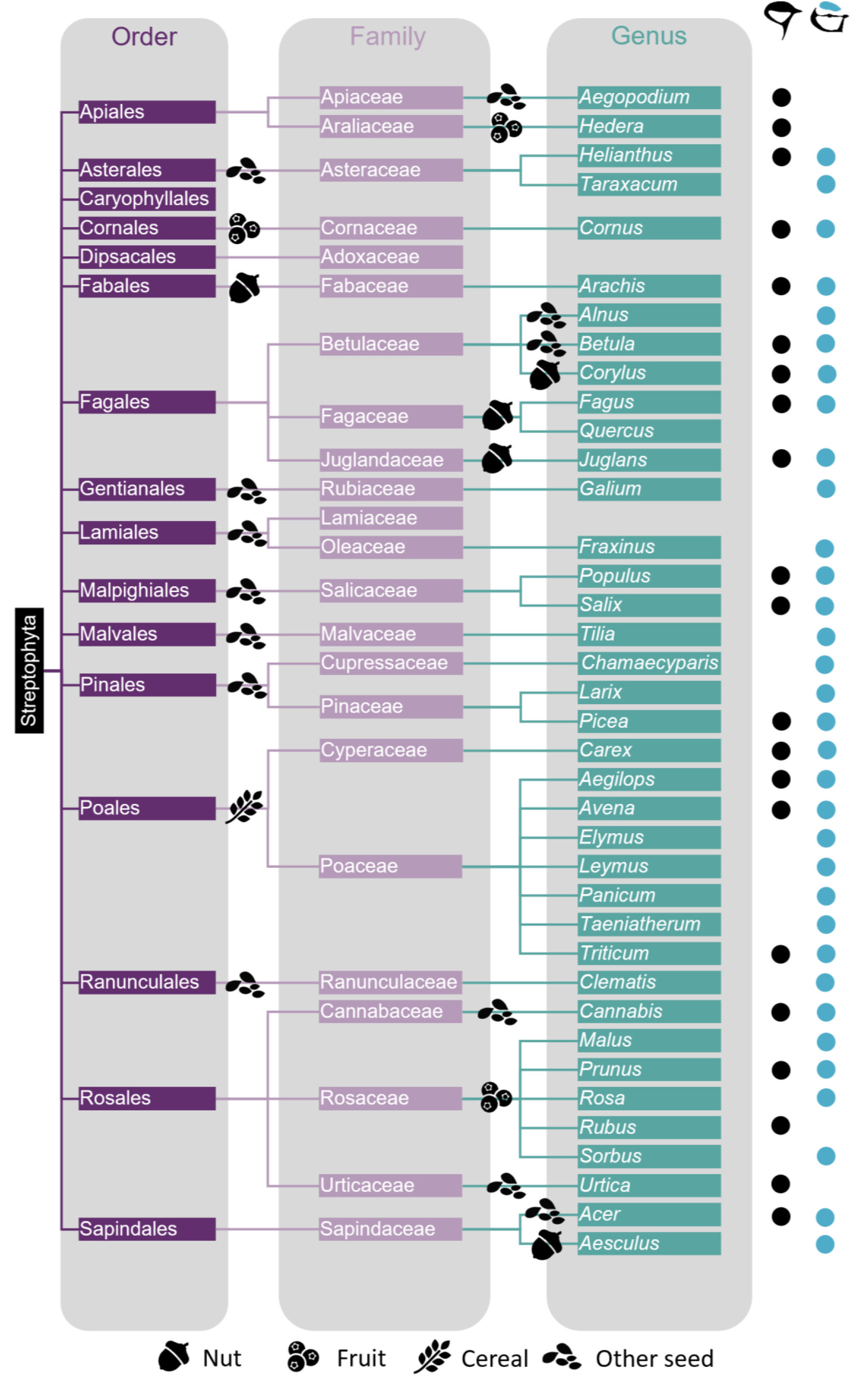
List of (a) arthropod and (b) plant taxa frequently detected in the droppings of great tits and blue tits. Genera included in the primary diet of great tits and blue tits are marked by a black and blue dot, respectively. Families that constitute the main diet of both species are shown in Fig. 1. All displayed orders were part of the main diet for both species, except for Gentianales, Malvales, and Psocoptera in great tits, and Apiales and Caryophyllales in blue tits. For plant taxa, symbols indicate whether the consumed item is expected to be a nut (i.e., hard-shelled seeds, usually high in fats and calories), a fruit (i.e., fleshy or soft plant parts, often high in sugars and vitamins), a cereal (i.e., grains from grass species, typically high in carbohydrates), or another type of seed (i.e., seeds from non-grass plants, including ornamental or wild species).

### 3.2. Diet taxonomic richness

In great tits, the total number of families detected per sample (i.e., per individual) decreased with increasing Tree cover (1000m) (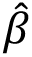=-0.063, χ²=6.55, P=0.010, R²m=0.076) (Fig. 2). The number of plant families detected per samples tended to depend on the interaction between ISA (1000m) and Month (χ²=10.54, P=0.061, R²m=0.058): In August, the number of plant families increased with increasing ISA (1000m) (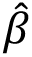=0.094, 95% CI [0.013 - 0.175]). Similar yet non-significant trends were observed in June and October (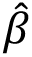=0.059, 95% CI [-0.012 - 0.129] and 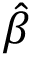=0.040, 95% CI [-0.010 - 0.090]) (Fig. 2). Results were similar in models using Tree cover (100m) and Tree cover (1000m) as explanatory variables. Moreover, the proportion of common families in each sample decreases with increasing ISA (100m) or ISA (1000m): The more urbanised the environment, the lower the proportion of common families compared to the average proportion for that month (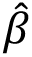=-0.003, χ²=7.53, P=0.006, R²m=0.034 and 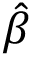=- 0.003, χ²=7.10, P=0.008, R²m=0.032) (Fig. 2). Similarly, the proportion of common plant and arthropod families in each sample decreases with increasing ISA (1000m) (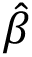=-0.002, χ²=5.91, P=0.008, R²m=0.040) (Fig. 2) and ISA (100m) (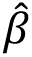=-0.003, χ²=6.28, P=0.012, R²m=0.062), respectively. Results were similar in models using Tree cover (100m) and Tree cover (1000m) as explanatory variables.

**Fig. 2:**
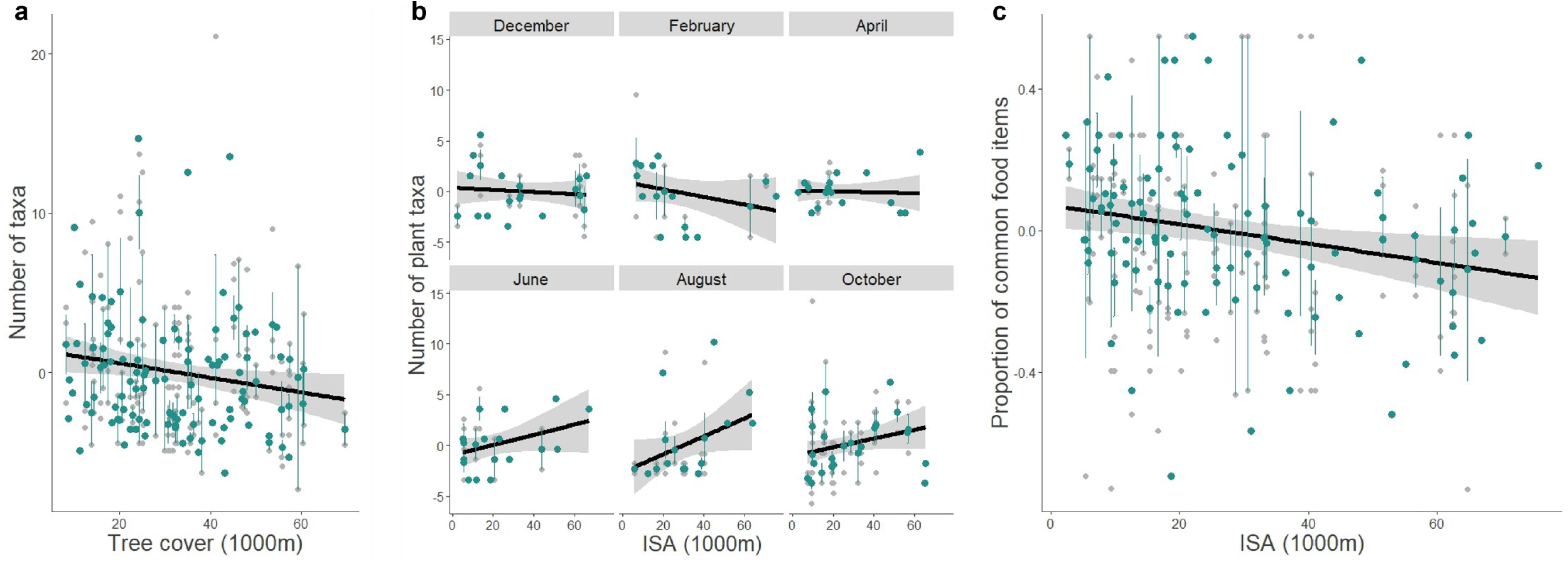
Variation in the number of (a) families and (b) plant families, and (c) in the proportion of common families (i.e., those occurring in at least 15% of individuals within a month) relative to the total number of families, in the diet of great tits along the rural-urban gradient (measured by either tree cover within a 1000m radius or imperviousness within a 1000m radius). The results for the number of plant families are presented by month. Individual values are shown in grey, while the mean ± standard error for each sampling site is shown in green.

Quite like the results on taxonomic richness at the family level, the total number of orders decreased with increasing Tree cover (1000m) (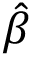=-0.043, χ²=5.09, P=0.024, R²m= 0.125). A similar trend was measured on plant taxonomic richness (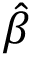=-0.026, χ²=3.67, P=0.055, R²m= 0.020). None of the variables were retained in the models on the proportion of common orders. In the same way, the total number of genera and the number of plant genera decreased with increasing Tree cover (1000m) (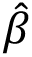=-0.048, χ²=4.22, P=0.040, R²m= 0.111 and 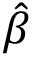=-0.040, χ²=3.32, P=0.025, R²m= 0.023, respectively). The proportion of common genera tended to decrease with increasing ISA (1000m) (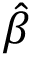=-0.002, F=2.89, P=0.091, R²= 0.009), while the proportion of common arthropod genera decreased with increasing ISA (100m) (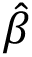=-0.001, F=4.08, P=0.045, R²=0.021).

None of the variables significantly explained taxonomic richness in blue tit diet.

### 3.3. Diet taxonomic composition

#### 3.3.1. Frequent food items in great tit and blue tit diet across the year

Both species exhibited noticeable seasonal shifts in their diet (Fig. 3). Asteraceae, Poaceae, Betulaceae, Fabaceae and Rosaceae were frequent food items throughout the year in both species; Pinaceae and Sapindaceae were also frequently detected during most of the year. As expected, the detection of Fabaceae, that corresponds exclusively to *Arachis sp.* (peanut) and that is present exclusively at bird feeders, was the highest from December to April in both species. The consumption of Fagaceae was extremely frequent in both species in October, during which it was detected in 65% of great tits 61% of blue tits. Its frequency in great tit diet remained rather high in December (46%) and February (30%). Blue tits consumed a larger range of arthropods, except during winter months when few or no arthropod were frequently detected. Philodromidae (running crab spiders) were frequently detected during most of the year, while Noctuidae (owlet moths), Cicadellidae (leafhoppers) and Aphididae (aphids) were frequently consumed from April to August, Theridiidae (cobweb spiders), Formicidae (ants), Curculionidae (weevils) and Clubionidae (sac spiders) from June to August, and Gracillaridae (leaf miner moths) and Cecidomyiidae (gall midges) from August to October. Fourteen other arthropod families were frequently but punctually (for one month only) detected between April and October. In the great tit, few arthropod groups were frequently detected over several months; those are Philodromidae from December to February, Anyphaenidae (ghost spiders) from February to June, and Curculionidae from June to August. Nine other arthropod families were frequently but punctually detected. The diversity of arthropod taxa in both species was the highest in June. Despite those general patterns, the frequency of several plant and arthropod taxa varied along the rural-urban gradient. Those relationships are detailed further below, separately for great tits and blue tits.

**Fig. 3:**
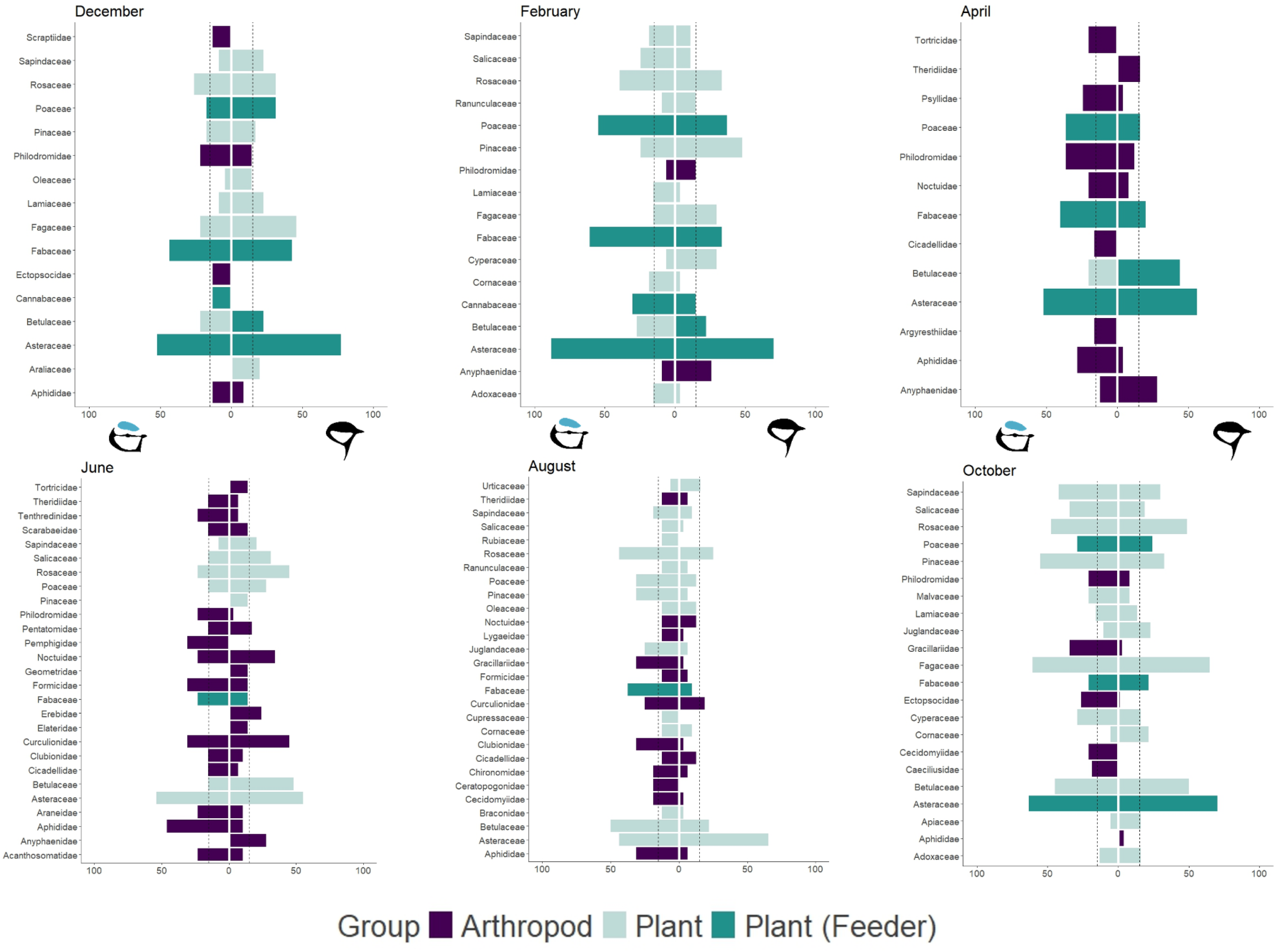
Food DNA detection frequency within great tit (on the right side) and blue tit (on the left side) diet across months. Arthropod and plant families are represented in green and violet, respectively, while plant families that were likely collected from bird feeders are represented in lavender. Fabaceae and Asteraceae are almost exclusively represented by *Arachis sp.* (peanuts; exclusively found at bird feeders) and *Helianthus sp.* (sunflowers), respectively. Betulaceae are represented by *Corylus sp.* (hazelnuts) as well as other taxa unlikely to be present at bird feeders; from December to in April, Betulaceae detected in great tits are mostly hazelnuts. Poaceae are represented by several cultivated plants *Triticum sp.* (wheat), *Avena sp.* (oats), *Zea mays* (corn), *Panicum miliaceum* (millet), *Secale sp.* (rye) as well as various grass taxa; from October to December, as well as in June, Poaceae mostly corresponds to cultivated plants. Only families present in at least 15% of the droppings/individuals of one of the two species sampled within the same month are displayed. Some taxa with a frequency slightly under 15% were selected due to rounding. The 15% threshold is visualized by a dashed line.

#### 3.3.2. Variations in the diet of great tits

Depending on the month and the taxonomic level considered, the level of urbanisation (ISA and/or Tree cover) significantly explained between 5.9% and 11.6% of the variation in great tit diet, while the habitat type significantly explained between 7.8% and 26.7% of the variation (Table 1). During the reproductive season (April-June), great tit diet was better explained by local scale urbanisation level (ISA or Tree cover in a 100m radius), while from October to February, great tit diet was better explained by landscape scale urbanisation level (ISA or Tree cover in a 1000m radius) and the habitat type. In August, great tit diet was not explained by the urbanisation level. Finally, urbanisation levels better explained the variation in diet composition at the family and genus level than at the order level.

**Table 1:** Results of the ordination analyses per month in (a) the great tit and (b) the blue tit. The table shows the results of the Monte Carlo permutation test (Anova), the variance explained by the selected constrained variables (i.e. the environmental variables that were the most important to explain the compositional changes, identified by forward selection), and the selected environmental variables. When several environmental variables were retained, we also indicate the percentage of variance explained by each of those variables. * although retained in the model, those variables do not significantly correlate to the RDA axis.

| (a) | Dec |  |  | Feb |  |  | Apr |  |  | Jun |  |  | Aug |  |  | Oct |  |  |
| --- | --- | --- | --- | --- | --- | --- | --- | --- | --- | --- | --- | --- | --- | --- | --- | --- | --- | --- |
|  | ORDER | FAMILY | GENUS | ORDER | FAMILY | GENUS | ORDER | FAMILY | GENUS | ORDER | FAMILY | GENUS | ORDER | FAMILY | GENUS | ORDER | FAMILY | GENUS |
| Anova | F=2.01,<br>P=0.002 | F=1.86,<br>P=0.001 | F=2.34,<br>P=0.001 | F=2.39,<br>P=0.033 | F=3.30,<br>P=0.001 | F=3.23,<br>P=0.002 | ∅ | F=2.95,<br>P=0.016 | F=5.02,<br>P=0.005 | F=1.76,<br>P=0.078 | F=2.67,<br>P=0.004 | F=2.34,<br>P=0.001 | ∅ | ∅ | ∅ | F=4.53,<br>P=0.001 | F=2.10,<br>P=0.001 | F=2.05,<br>P=0.001 |
| Variance explained | 23.3% | 23.3% | 28.7% | 8.7% | 11.6% | 11.4% |  | 11.4% | 25.2% | 6.1% | 9.0% | 22.2% |  |  |  | 5.9% | 15.6% | 13.1% |
| ISA (100m) |  |  |  |  |  |  |  |  |  |  |  | 11.2% |  |  |  |  | 3.5% |  |
| ISA (1000m) |  |  | 6.0% |  |  |  |  |  |  |  |  |  |  |  |  |  |  |  |
| Tree cover (100m) |  |  |  |  |  |  |  |  |  |  |  | 5.6%* |  |  |  |  |  |  |
| Tree cover (1000m) |  |  |  |  |  |  |  |  |  |  |  | 5.4%* |  |  |  |  | 4.2% | 5.3% |
| Habitat |  |  | 22.7% |  |  |  |  |  |  |  |  |  |  |  |  |  | 8.2% | 7.8% |

| (b) | Dec |  |  | Feb |  |  | Apr |  |  | Jun |  |  | Aug |  |  | Oct |  |  |
| --- | --- | --- | --- | --- | --- | --- | --- | --- | --- | --- | --- | --- | --- | --- | --- | --- | --- | --- |
|  | ORDER | FAMILY | GENUS | ORDER | FAMILY | GENUS | ORDER | FAMILY | GENUS | ORDER | FAMILY | GENUS | ORDER | FAMILY | GENUS | ORDER | FAMILY | GENUS |
| Anova | ∅ | ∅ | ∅ | F=2.71,<br>P=0.006 | F=2.42,<br>P=0.005 | F=1.79,<br>P=0.052 | F=1.72,<br>P=0.008 | F=4.23,<br>P=0.001 | F=4.50,<br>P=0.001 | ∅ | ∅ | ∅ | ∅ | F=1.63,<br>P=0.043 | F=1.98,<br>P=0.012 | ∅ | F=1.70,<br>P=0.001 | F=1.92,<br>P=0.001 |
| Variance explained |  |  |  | 8.1% | 7.2% | 5.5% | 25.7% | 15.5% | 16.4% |  |  |  |  | 10.4% | 12.4% |  | 21.0% | 23.1% |
| ISA (100m) |  |  |  |  |  |  |  |  |  |  |  |  |  |  |  |  |  |  |
| ISA (1000m) |  |  |  |  |  |  |  |  |  |  |  |  |  |  |  |  | 5.0% | 5.7% |
| Tree cover (100m) |  |  |  |  |  |  |  |  |  |  |  |  |  |  |  |  |  |  |
| Tree cover (1000m) |  |  |  |  |  |  |  |  |  |  |  |  |  |  |  |  |  |  |
| Habitat |  |  |  |  |  |  |  |  |  |  |  |  |  |  |  |  | 15.0% | 17.4% |

The variation in the frequency of food items in great tit diet in response to urbanisation depended on both the taxon and the month. Some taxa showed a consistent decline in frequency as urbanisation increased, while others exhibited an increase. For certain taxa, this relationship changed across different months (Fig. 4). Interestingly, arthropod consumption during the breeding season showed variation in response to urbanisation levels. In April, the frequency of Anyphaenidae ghost spiders (specifically from the *Anyphaena* genus) decreased as urbanisation increased. In June, the consumption of Lepidoptera, particularly from the Erebidae moth family, and Coleoptera, especially the *Phyllobius* and *Polydrusus* weevil genera, also declined with higher urbanisation levels, whereas the consumption of Formicidae increased; the frequency of Asteraceae (specifically from sunflowers, *Helianthus*) also increased with increasing urbanisation level during this season while the one of Betulaceae (specifically from hazelnuts, *Corylus*) decreased (in April only). Another notable change along the rural-urban gradient was the increased consumption frequency of several plant taxa, specifically Sapindales (mainly various maple species, *Acer*), Cornales (primarily dogwoods, *Cornus*), Juglandaceae (mostly walnut trees, *Juglans*), and Fagaceae (mainly beech trees, *Fagus*) during the month of October, but the decreased frequency of *Helianthus* and Fabales (exclusively of *Arachis*). The higher frequency of *Fagus* in more urbanised areas continued through December and February. Finally, during the winter month of February, the frequency of Fabales/Fabaceae (exclusively represented by *Arachis*) increased with increasing urbanisation level, while the frequency of Betulaceae, especially of *Corylus*, tended to decrease. Spiders were the only arthropods commonly detected during this month, but urbanisation was related to a shift in the taxa that are consumed. More specifically, the frequency of Anyphaenidae decreased while the frequency of Philodromidae tended to increase along the rural-urban gradient. In addition, the frequency of Pinales (which exclusively comprises the *Picea* genus) and Poales, especially of Cyperaceae (which exclusively comprises *Carex duriuscula*) decreased, while the frequency of Fagaceae, especially of *Fagus*, increased along the rural-urban gradient.

**Fig. 4:**
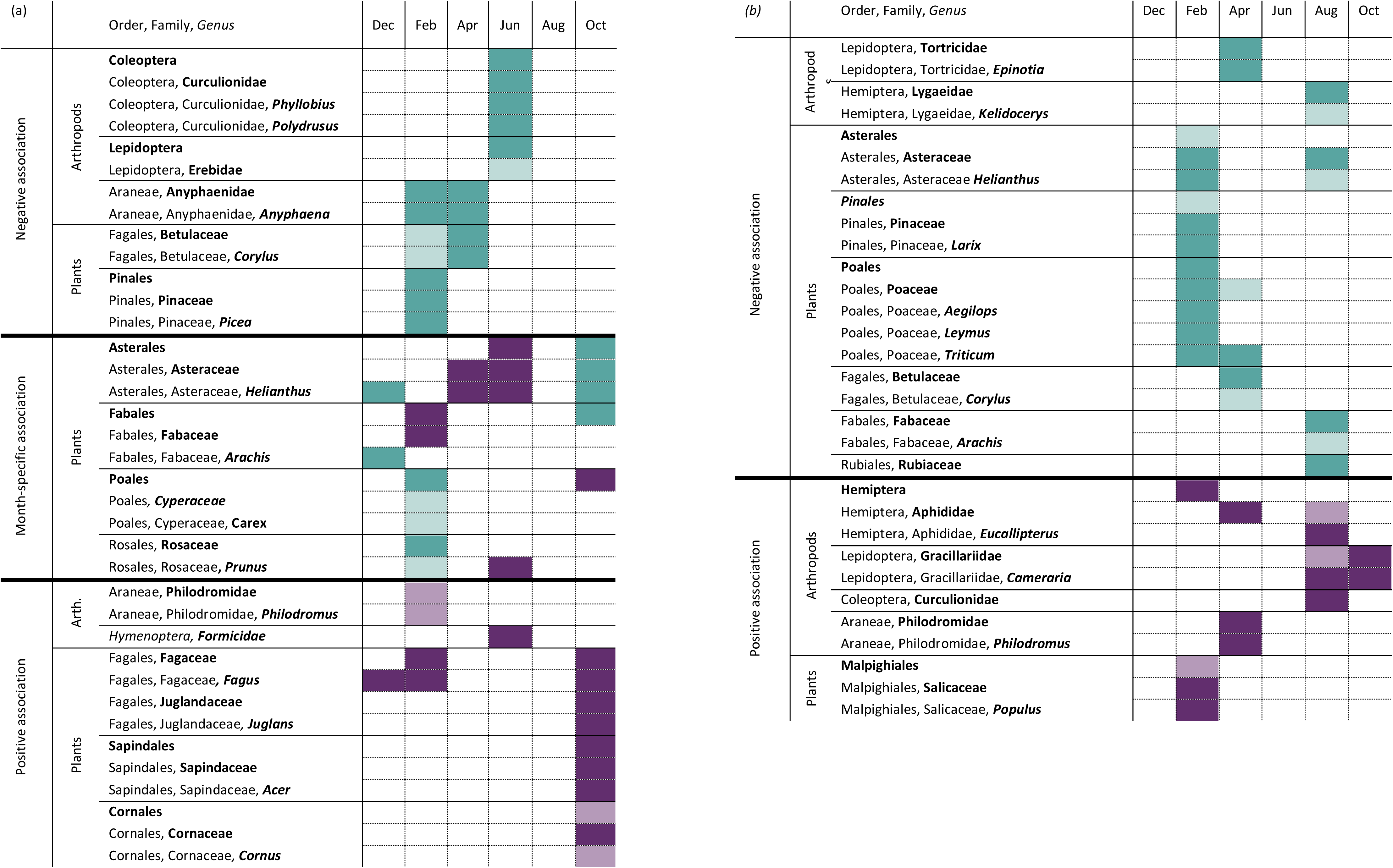
Main taxonomic groups explaining variations in (a) great tit and (b) blue tit diet composition along the rural-urban gradient. Groups which presence in the diet decreased and increased along the rural-urban gradient are highlighted in teal and purple, respectively. This summary is based on the interpretation of the ordination analysis for each month. Only groups contributing to 10% or more of the RDA components are shown. Groups contributing between 5% and 10% are also shown (in a lighter colour) when their association aligns with patterns observed at a higher or lower taxonomic level, or during the preceding or following sampling month. Results in December and October in great tits, and October in blue tits are also driven by habitat differences (see Fig. 5).

In December and October, great tit diet was also explained by the habitat type. In December, RDA1 significantly explained 19.1% of the variation in diet composition at the genus level (F=7.76, P=0.001). It mainly correlated to the habitat (r=0.91) and, in a lower extent, to ISA (1000m) (r=0.57). It separates forest remnants and residential areas, characterised by a higher frequency of *Helianthus* and *Arachis*, from parks and garden squares, characterised by a higher frequency of *Fagus* (Fig. 5). Similar results were obtained at the order and family levels, except that the frequencies of Poales and Poaceae (from which 78% of the detections belongs to the cultivated plants *Triticum*, *Avena*, *Zea.*, *Panicum* and *Secales*) varied similarly to those of Asterales/Asteraceae and Fabales/Fabaceae. The effect of the habitat could not be tested during February, the other winter month in our dataset, because of the underrepresentation of some habitats. However, the variation in the relative frequency of Fabaceae indicates a different pattern compared to that observed in December, with relatively high frequencies in garden squares. In October, RDA1 explained 7.2% of the variation in diet composition at the genus level (F=5.60, P=0.001). It mainly correlates with Tree cover (1000m) (r=-0.83) and, in a lower extent, to the habitat (r=0.60). It separates forests, characterised by a higher frequency of *Helianthus*, from garden squares and forest remnants, characterised by a higher frequency of *Fagus*, *Acer, Juglans* and *Cornus* (Fig. 5). The results were similar at the family level.

**Fig. 5:**
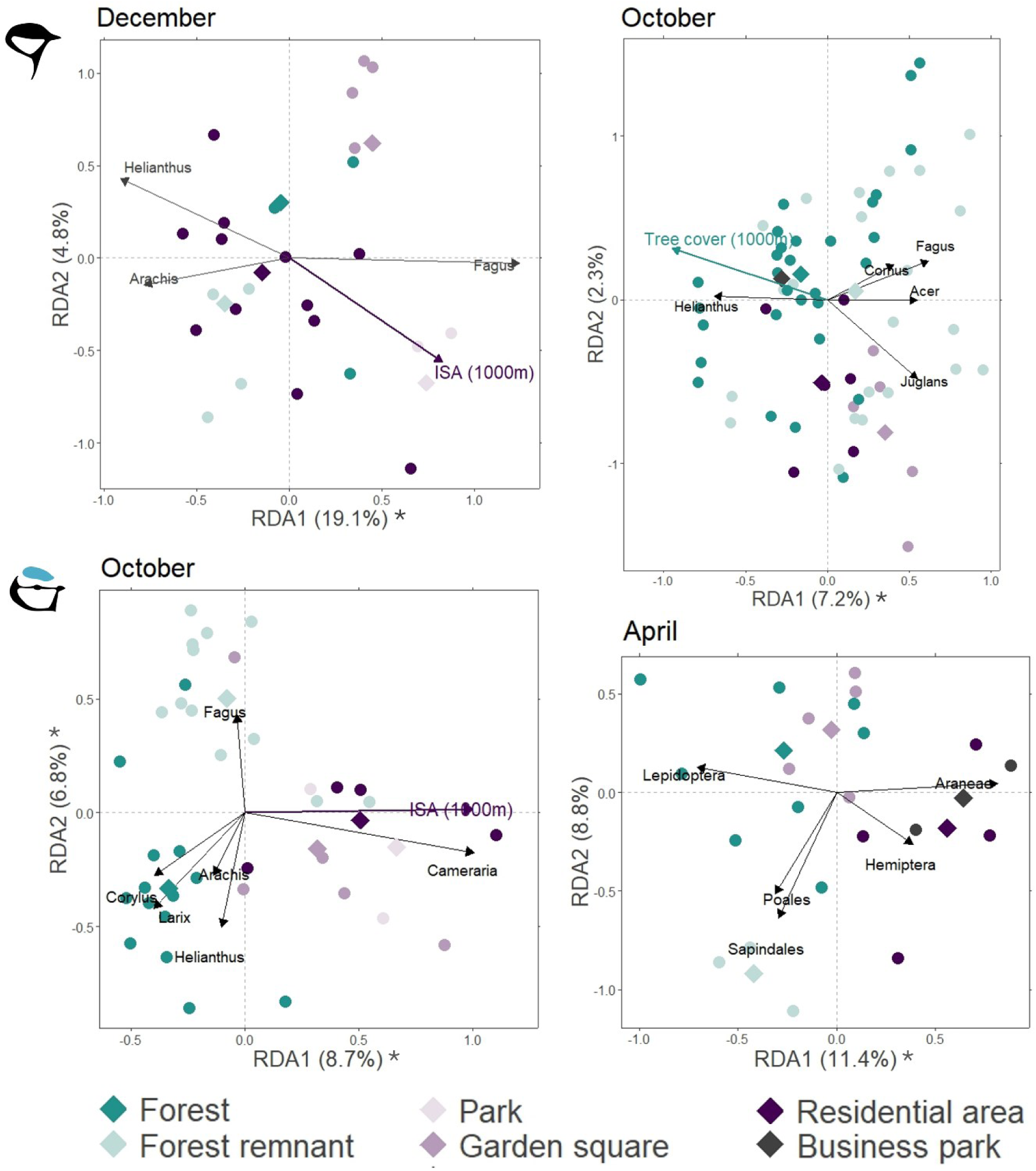
Redundancy analysis triplots in the great tit (top panel) and the blue tit (bottom panel). Different symbol colours highlight clusters by habitat type. The diamonds show the centroid for each habitat. The main taxonomic groups explaining the ordination are highlighted by black vectors (they explain more than 10% of the ordination). The continuous environmental variables significantly explaining differences in diet composition between individual are represented by coloured vectors. Asterisks indicate significant axes.

#### 3.3.3. Variations in the diet of blue tits

Depending on the month and the taxonomic level considered, the level of urbanisation (ISA and/or Tree cover) significantly explained between 5% and 16.4% of the variation in blue tit diet, while the habitat type significantly explained between 15% and 25.7% of the variation (Table 1). During August and October, blue tit diet was better explained by landscape-scale urbanisation level (ISA 1000m radius); the habitat type also significantly explained diet variations in October. Contrary to what was measured in the great tit, blue tit diet during April was associated with local-scale and landscape-scale urbanisation levels, as well as by the habitat type. In February, blue tit diet was significantly associated with local-scale tree cover. Blue tit diet during June and December was not explained by the urbanisation level, nor by the habitat. In June, it is likely that our model lacked statistical power due to low sample size. Finally, except for the month of February, urbanisation level only explained diet at the family and genus levels.

Like for great tits, the variation in the frequency of food items in blue tit diet in response to urbanisation depended on both the taxon and the month (Fig. 4). Interestingly, in April (during the breeding season), the frequency of Tortricidae moths, especially from the genus *Epinotia*, decreased, while the frequency of Philodromidae crab spiders, especially from the genus *Philodromus*, and of Aphididae, increased with increasing urbanisation level. The urbanisation level was also associated to changes in arthropod consumption in August, with birds in more urbanised areas consuming more frequently Gracillaridae moths, almost exclusively of the horse-chestnut leaf miner *Cameraria orhidella*, Curculionidae (weevils) and Aphididae, especially of the linden aphid *Eucallipterus tiliae*, while they consumed less frequently Lygaeidae true bugs, especially of the *Kleidocerys* seed bugs. The positive association with Gracillaridae carried on until October. Finally, the consumption of food items likely collected from bird feeders (*Helianthus*, *Triticum*, *Corylus* and *Arachis*) was negatively associated with urbanisation in February, April and August.

In April and in October, blue tit diet also varied between habitats. In April, RDA1 explained 11.5% of the variation in diet composition at the order level (F=3.06, P=0.051). It separates forest remnants and forests, characterised by a higher frequency of Lepidoptera, and in a lower extent of Poales and Sapindales, from business parks and residential areas, characterised by a higher frequency of Aranea and, in a lower extent, of Hemiptera (Fig. 5). In October, RDA1 and RDA2 significantly explained 8.7% (F=3.64, P=0.003) and 6.8% (F=2.82, P=0.009) of the variation in diet composition at the genus level, respectively. RDA1 correlated to both ISA (1000m) (r=0.97) and the habitat (r=0.94); it separates forests from parks, garden squares and residential areas, characterised by a higher frequency of *Cameraria* moths. The same result was obtained at the family level Gracillariidae. RDA2 correlated only to the habitat (r=0.96); it separates forests, parks and garden squares, characterised by a higher frequency of *Helianthus*, *Picea*, *Larix*, *Corylus* and *Arachis*, from forest remnants, characterised by a higher frequency of *Fagus* (Fig. 5).

### 3.4. Arthropod consumption – arthropod availability match

Significant positive correlations between the consumption of arthropod and their abundance in trees and bushes were detected for few taxonomic groups. In great tits, the consumption of Anyphaenidae (ghost spiders) in February increased with their abundance in trees and bushes (r=0.49, t=2.82, P=0.009 and r=0.55, t=3.33, P=0.003), while the consumption of Theridiidae (tangle-web spiders) in April and of Curculionidae (true weevils) in June increased with their respective abundances in bushes (r=0.53, t=3.29, P=0.003 and r=0.41, t=2.16, P=0.041, respectively). In blue tits, the consumption of Cicadellidae (leafhoppers) in June increased with their abundance in tree canopy (r=0.66, t=2.89, P=0.015), while the consumption of Ectopsocidae (bark lice) in October and Philodromidae (crab spiders) in December increased with their respective abundances in bushes (r=0.47, t=3.12, P=0.004 and r=0.42, t=2.15, P=0.044, respectively).

## 4. Discussion

Our study showed that urbanisation alters the diet diversity and composition of both great tits and blue tits. In great tits, urban areas encourage a broader diet, and in particular increases the consumption of plant-based food items such as seeds and fruits. This increased dietary diversity likely stems from the varied plant species found in urban green spaces. Maples, walnuts, and other ornamental tree species are abundant in urban settings, and their presence was reflected in the diets of great tits. This was particularly evident from June to October, when the availability of fruits, seeds, and nuts from these trees is at its peak. In addition, our results show that the consumption of rare taxa (i.e., taxa that have been detected in one or a few individuals only) increases along the rural-urban gradient whatever the season. Importantly, this relationship was similar when considering plant and arthropod taxa separately. Urban environments, therefore, appear to promote an opportunistic feeding behaviour in great tits. This aligns with the idea that urban environments are composed of small patches of diverse habitats, leading to increased variability between individuals within urban populations compared to rural ones (Thompson et al., 2022). In contrast, blue tits did not show a significant change in diet diversity along the rural-urban gradient. This suggests that blue tits maintain a broad and flexible diet regardless of the habitat, incorporating both arthropods and plant items in urban and rural areas alike. As a matter of fact, 33 plant and 26 arthropod genera were frequently detected as food sources in this species, while only 21 plant and 8 arthropod genera were regularly detected in the great tit.

Seasonal differences in diet composition revealed that both species rely heavily on arthropods during the breeding season (April to June), although urbanisation appears to reduce the availability of key arthropod groups, particularly for great tits. Our findings align with previous research, indicating that urban great tits and blue tits experience a reduction in essential arthropods such as caterpillars during this critical time (Marciniak et al., 2007; Nadolski et al., 2021; Piano et al., 2020; Seress et al., 2018). Great tits in urban environments also showed reduced consumption of spiders and weevils but higher consumption of ants. In blue tits, the less frequent consumption of caterpillars was associated to a higher consumption of other arthropod groups, specifically spiders and aphids. Because their consumption was correlated to their abundance in the environment or because the abundance of those groups varied similarly along the rural-urban gradient, our results suggest that shifts in arthropod consumption reflect their availability in the environment (Chatelain et al., 2023).

Egg formation requires essential nutrients for the shell, yolk, and albumen, with protein-rich arthropods such as caterpillars and spiders providing essential amino acids for embryo development (Willems et al., 2014). A lower intake of protein-rich arthropods as for example caterpillars and spiders could limit the birds’ ability to produce eggs and/or to provide essential amino acids for embryo development (Ramsay & Houston, 1998). Spiders also supply calcium, crucial for strong eggshells (Graveland & Gijzen, 2002); reduced spider intake can lead to weaker shells and lower hatching success (Graveland & Drent, 1997). In urban settings, however, great tits consumed more sunflower seeds, which are high in omega-6 fatty acids but lack the proteins and omega-3 fatty acids found in arthropods (Andersson et al., 2015). An imbalance favouring omega-6 over omega-3 can lead to chronic inflammation, reduced immunity, and compromised egg quality, potentially resulting in lower hatching success and chick survival (Attia et al., 2022; Gutiérrez et al., 2019). In urban areas, both great tits and blue tits showed decreased moth consumption, suggesting a deficiency in omega-3 intake, though blue tits may mitigate this by consuming more spiders and aphids. Further research is needed on the nutritional content of urban diets and their implications for bird fitness to understand the full impact of these dietary shifts on reproductive success.

From October to February (i.e., outside of the breeding season), the diet of great tits was influenced by landscape-scale urbanisation (within a 1000 m radius) and habitat type, reflecting the birds’ expanded winter foraging ranges when territorial behaviours decrease (Cox & Gaston, 2018; Nakamura & Shindo, 2001). In October, great tits in urban areas seem to have benefited from a greater variety of available nuts, seeds, and berries, especially in garden squares and forest remnants where ornamental plants are common. In contrast, managed forested areas tend to consist of a few dominant tree species, such as Norway spruce and European larch, which are reflected in the diet through increased consumption of these seeds in less urbanised areas. The diet of blue tits from August to October was also shaped by landscape-scale urbanisation and habitat type, suggesting that they utilize a broad foraging range outside of the breeding season. However, in February, blue tit diet was more closely associated with local tree cover, which may indicate a focus on foraging efficiency during the coldest winter month by staying near food-rich areas as for example tree cover or feeders. Contrary to great tits, blue tits outside the reproductive season frequently consumed various arthropod prey, such as spiders, moths, weevils, aphids, as well as flies and barklice. Urban blue tit populations showed increased consumption of aphids, leaf miner moths, and weevils, likely due to their prevalence in ornamental urban trees such as lindens and horse chestnuts. These findings highlight the importance of considering habitat diversity along the rural-urban gradient when studying urban ecosystems. Indeed, habitat diversity significantly influences vegetation composition and environmental factors such as the availability of anthropogenic food, as well as noise, light, and chemical pollution. These factors, which are not captured by measurements of land cover, play a crucial role in shaping urban environments and need more attention in future work to unravel the consequences of urbanization on animals.

Outside the breeding season, diet may impact bird health as well as the expression of secondary sexual traits (e.g., carotenoid-based plumage colouration and melanic breast stripe), which are essential for survival and future reproductive success (Meunier et al., 2011; Weaver et al., 2018). Urbanisation has been associated with reduced carotenoid deposition in feathers during moult (from August to October)—a phenomenon known as urban dullness (Leveau, 2021). Carotenoids are obtained from fruits (e.g., rosehips, dogwood berries, elderberries, apples, rowan berries, brambles), seeds (sunflower, pumpkin, niger seeds) and arthropods (mainly caterpillars feeding on yellow or orange flowers, fruits, or leafy greens). However, both bird species frequently consumed fruits such as plums, rosehips, and brambles in August and October, with no significant difference in their consumption frequency along the rural-urban gradient. This finding, along with the observed urban dullness phenomenon in the populations we studied (unpublished data), suggests that the carotenoid content in similar food items may vary across the rural-urban gradient (Isaksson, 2009; Isaksson & Andersson, 2007), that the phenomenon may result from a trade-off in carotenoid allocation (Koch & Hill, 2018) or that the deposition of carotenoids in the feathers does not strictly depend on the diet during moult. Investigating the impact of diet on carotenoid-based coloration will be a key step in understanding how urbanization affects the production of secondary sexual traits and reproductive success.

Additionally, beech mast is a critical winter food source for both species, especially for juvenile great tits (Kallander, 1981; Perdeck et al., 2000; Woodman et al., 2024). The increased beech nut consumption along the rural-urban gradient in forest remnants highlights the essential role of forest remnants in supporting urban bird populations by supplying native food sources crucial for survival. Similar patterns are likely to be observed across Europe, where beech (*Fagus sylvatica*) is a dominant species in temperate forests. This highlights the need for further research specifically examining the role of forest remnants in enhancing resource availability and supporting bird populations in urbanised landscapes.

Finally, our study revealed that bird feeding plays a significant role in the diets of great tits and blue tits, with both species consuming high-fat foods such as sunflower seeds and peanuts from October to April. The presence of peanuts in their diet year-round also suggests that some households provide food for birds throughout the entire year. This consumption varied by habitat and urbanisation level, indicating that bird feeding practices differ along the rural-urban gradient. Great tits, particularly in residential and forest remnant areas where feeders are more common, consumed more sunflower seeds and peanuts in winter months. Interestingly, the reduced consumption of feeder items in October in more urbanised areas might indicate that birds prefer to forage from naturally available seeds and berries than to visit bird feeders, where competition (Davis, 2009; Francis et al., 2018; Miller et al., 2017), predation risk (Dunn & Tessaglia, 1994) and stress related to human proximity may be higher (Davis, 2009; Dunn & Tessaglia, 1994; Miller et al., 2017; Tryjanowski et al., 2016). In contrast to great tits, blue tits showed either stable or decreased consumption of feeder-provided food along the rural-urban gradient. Previous studies support this pattern, suggesting that blue tits are more abundant at rural feeders (Tryjanowski et al., 2015). Blue tits, being lower in dominance at bird feeders, may face competition from species such as great tits, chaffinches, and house sparrows in urban areas, limiting their access to feeders (Francis et al., 2018). Alternatively, the broader diet of blue tits compared to great tits may allow them to forage on a variety of natural food sources, including arthropods, even in urban settings. Bird feeding has been linked to improved winter survival (Shutt et al., 2021), higher reproductive success (Robb et al., 2008; Ruffino et al., 2014), and even increased population trends (Plummer et al., 2019; Shutt et al., 2021). The frequent consumption of high-fat foods, such as sunflower seeds and peanuts, during winter likely supports the survival of the populations we studied. However, our findings suggest that birds forage from bird feeders nearly year-round, which could impact long-term fitness by altering the intake of essential amino acids and micronutrients crucial for immune function, feather quality, and reproduction. This result highlights the importance of understanding the role that bird feeding may play in the lower productivity of urban populations compared to their rural counterparts.

In conclusion, our results address an important gap in understanding the causal link between urbanisation and changes in animal fitness. The current findings suggest that urbanisation alters the availability of preferred food items, forcing birds to adjust their diet. For food generalists such as great tits, this leads to more opportunistic foraging, with an increased reliance on energy-dense foods as for example sunflower seeds and a reduced intake of nutrient-rich arthropods during the reproductive season. For blue tits, largely insectivorous, urbanisation causes a shift in the types of food consumed, particularly in the arthropod prey they rely on. Our findings also emphasize the need for urban ecology studies to consider both urbanisation level and habitat type, as habitat variation influences plant assemblages, arthropod prey availability and human activities, all of which impacting bird diets, fitness and ultimately the eco-evolution of birds in an urbanizing world.

## 5 Acknowledgments

We would like to thank Angelika Fritz for her assistance with field data collection and DNA extractions, and Samuel Caro for providing samples for the optimisation of the metabarcoding protocols. We also extend our gratitude to Christiane Winter for providing training and guidance throughout every step of the molecular work conducted in this study. This study was supported by the Austrian Science Fund (doi: 10.55776/M2628). The funder had no role in study design, data collection and analysis, the decision to publish, or in preparation of the paper. The project was approved by the Austrian Ornithological Center, the Austrian Federal Ministry of Education, Science and Research and the Office of the Tyrolean Provincial Government. The authors declare no conflict of interest.

