## Supplementary material for "Urbanisation and Habitat Shape Resource-Driven Dietary Shifts in Wild Birds"

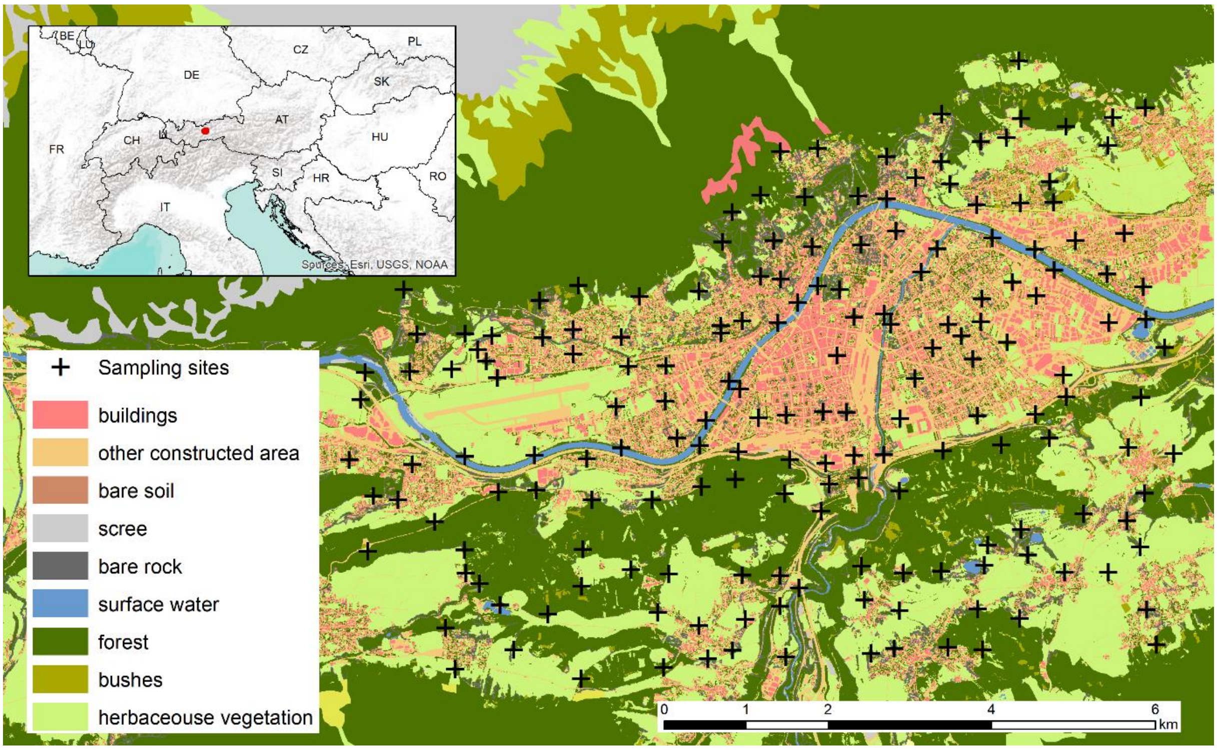


**Fig. 1. Location of Innsbruck within Europe and overview of the 180 sites in Innsbruck and its surroundings where great tits and blue tits were sampled from October 2020 to August 2021.** The map highlights the land cover in the sampling area extracted from the Land Information System Austria.

**Table 1. Number of individuals per month and habitat type in great tits and blue tits.**

| | Forest | Forest remnant | Residential area | Garden square | Park | Business park | Total |
|---|---|---|---|---|---|---|---|
| **December** | 5\|3 | 5\|1 | 17\|14 | 5\|4 | 3\|1 | 0\|0 | 35\|23 |
| **February** | 14\|11 | 3\|1 | 3\|9 | 6\|7 | 1\|3 | 0\|2 | 27\|33 |
| **April** | 14\|10 | 4\|3 | 3\|5 | 3\|5 | 0\|0 | 1\|2 | 25\|25 |
| **June** | 20\|11 | 2\|0 | 0\|0 | 7\|2 | 0\|0 | 0\|0 | 29\|13 |
| **August** | 23\|10 | 1\|0 | 2\|3 | 6\|3 | 0\|0 | 0\|0 | 32\|16 |
| **October** | 35\|14 | 23\|12 | 6\|4 | 5\|5 | 0\|3 | 1\|0 | 74\|38 |
| **Total** | **111\|59** | **38\|13** | **31\|34** | **32\|25** | **4\|6** | **2\|4** | **222\|148** |

**Table 2. List of genera likely to occur as pollen each month.** The number of occurrences removed from the dataset is indicated in brackets.

| Month | Genus |
|---|---|
| February | Alnus (10) |
| April | Acer (1), Aegilops (0), Aegopodium (5), Betula (0), Cornus (0), Fagus (3), Juglans (4), Larix (6), Picea (0), Pinus (6), Prunus (0), Quercus (5), Salix (0) |
| June | Aegilops (6), Aegopodium (2), Aesculus (2), Carex (0), Clematis (0), Cornus (0), Elymus (2), Fagus (4), Leymus (2), Picea (0), Sambucus (5), Tilia (4) |
| October | Hedera (12) |

**Table 3. Number of taxa detected per month and per species.** The last column displays the number of unique taxa detected across the year.

| Month | | Dec | Feb | Apr | Jun | Aug | Oct | TOTAL | |
|---|---|---|---|---|---|---|---|---|---|
| No. genus | GT | 100 | 85 | 77 | 122 | 136 | 179 | | |
| | BT | 92 | 107 | 97 | 74 | 86 | 166 | | |
| No. genus (excluding pollen) | GT | 100 (31) | 84 (31) | 70 (39) | 115 (73) | 136 (76) | 178 (67) | 392 (233) | 547 (351) |
| | BT | 92 (36) | 106 (37) | 92 (55) | 69 (46) | 86 (46) | 165 (89) | 335 (201) | |
| No. abundant genus | GT | 10 (1) | 13 (2) | 4 (1) | 15 (6) | 5 (1) | 14 (0) | 29 (8) | 67 (30) |
| | BT | 10 (2) | 14 (0 | 10 (5) | 16 (11) | 29 (7) | 18 (5) | 59 (26) | |
| No. families | GT | 53 | 54 | 53 | 68 | 80 | 90 | | |
| | BT | 59 | 63 | 67 | 37 | 55 | 94 | | |
| No. families (excluding pollen) | GT | 53 (25) | 54 (26) | 50 (31) | 66 (41) | 80 (50) | 89 (42) | 154 (99) | 199 (135) |
| | BT | 59 (29) | 63 (32) | 64 (40) | 35 (25) | 55 (35) | 93 (58) | 148 (97) | |
| No. abundant families | GT | 12 (1) | 12 (2) | 6 (2) | 18 (10) | 5 (1) | 14 (0) | 31 (12) | 53 (30) |
| | BT | 12 (4) | 13 (0) | 11 (7) | 20 (14) | 27 (13) | 18 (6) | 46 (25) | |
| No. orders | GT | 28 | 29 | 20 | 27 | 30 | 41 | | |
| | BT | 30 | 28 | 29 | 19 | 26 | 34 | | |
| No. orders (excluding pollen) | GT | 28 (10) | 29 (10) | 18 (7) | 25 (7) | 30 (8) | 40 (12) | 46 (16) | 51 (19) |
| | BT | 30 (11) | 28 (10) | 27 (12) | 17 (8) | 26 (10) | 33 (10) | 42 (16) | |
| No. abundant orders | GT | 12 (2) | 11 (2) | 10 (6) | 15 (6) | 11 (6) | 12 (1) | 20 (6) | 25 (7) |
| | BT | 15 (5) | 16 (5) | 11 (6) | 11 (5) | 19 (6) | 17 (7) | 22 (7) | |
